# Functional cortical localization of the tongue using corticokinematic coherence with a deep learning-assisted motion capture system

**DOI:** 10.1101/2021.08.18.456754

**Authors:** Hitoshi Maezawa, Momoka Fujimoto, Yutaka Hata, Masao Matsuhashi, Hiroaki Hashimoto, Hideki Kashioka, Toshio Yanagida, Masayuki Hirata

## Abstract

Measuring the corticokinematic coherence (CKC) between magnetoencephalographic and movement signals using an accelerometer can evaluate the functional localization of the primary sensorimotor cortex (SM1) of the upper limbs. However, it is difficult to determine the tongue CKC because an accelerometer yields excessive magnetic artifacts. We introduce and validate a novel approach for measuring the tongue CKC using a deep learning-assisted motion capture system with videography, and compare it with an accelerometer in a control task measuring finger movement. Twelve healthy volunteers performed rhythmical side-to-side tongue movements in the whole-head magnetoencephalographic system, which were simultaneously recorded using a video camera and examined offline using a deep learning-assisted motion capture system. In the control task, right finger CKC measurements were simultaneously evaluated via motion capture and an accelerometer. The right finger CKC with motion capture was significant at the movement frequency peaks or its harmonics over the contralateral hemisphere; the motion-captured CKC was 84.9% similar to that with the accelerometer. The tongue CKC was significant at the movement frequency peaks or its harmonics over both hemispheres, with no difference between the left and right hemispheres. The CKC sources of the tongue were considerably lateral and inferior to those of the finger. Thus, the CKC based on deep learning-assisted motion capture can evaluate the functional localization of the tongue SM1. In this approach, because no devices are placed on the tongue, magnetic noise, disturbances due to tongue movements, risk of aspiration of the device, and risk of infection to the experimenter are eliminated.

## 1. Introduction

The tongue plays an important role in various critical human functions, including swallowing, mastication, and speech, and can perform sophisticated movements. The area of the primary sensorimotor cortex (SM1) representing the tongue occupies a wide distribution relative to its actual size in the body (Penfield and Boldrey, 1937), suggesting the functional importance of the SM1 of the tongue region. However, as it is difficult to measure electromagnetic cortical signals during tongue movements without artifact contamination because of the short distance between the tongue and brain, the cortical representation of the tongue regions has rarely been examined. Thus, it is important to establish robust methods for evaluating the functional localization of the tongue region to reveal the central mechanisms of fine tongue movements. Moreover, as the tongue is innervated by both hypoglossal nerves from the bilateral SM1 (Penfield and Boldrey, 1937). Therefore, neurosurgical operation of the target side of the tongue SM1 does not generally deteriorate the tongue motor functions compared with hand motor dysfunctions that frequently occur when operating on the hand SM1. However, it is essential to evaluate the presurgical localization of the SM1 of the tongue region to minimize the deterioration of tongue motor functions after surgery, which would significantly reduce the post-surgery quality of life, since dysfunctions in critical tongue motor functions can potentially cause dysphagia and silent aspiration (Meadows, 1973; Horner and Massey, 1988; Robbins et al., 1993). Thus, during neurosurgery, it is critical to evaluate the somatotopic source localization of the tongue region before the surgery.

Corticokinematic coherence (CKC) is a useful approach for identifying the SM1 of fingers in healthy adults (Bourguignon et al., 2011, 2019), newborns (Smeds et al., 2017), and patients with impaired spinocortical proprioceptive pathways in Friedreich ataxia (Marty et al., 2019). Conventional CKC methods quantify the coupling between magnetoencephalographic (MEG) signals and finger kinematics, which are measured using an accelerometer (ACC) during repetitive, rhythmic, and voluntary finger movements (Bourguignon et al., 2011; 2013). Previous studies have shown that the CKC mainly reflects the proprioceptive input into the SM1 (Piitulainen et al., 2013a; Bourguignon et al., 2015; 2017); this feature is comparable to the strongest deflections observed in the cortical movement evoked fields (MEFs) associated with voluntary finger movements (Cheyne et al., 1989; 1997; Gerloff et al., 1998). However, it is difficult to apply this technique to regions of the tongue using an ACC because the ACC produces excessive magnetic artifacts, which easily contaminate the cortical magnetic activity due to the short distance between the tongue and MEG sensors. It is also technically challenging to set an ACC on narrow and wet tongue regions. Moreover, ACCs with cables have the disadvantage of sometimes disturbing the smooth movements of the tongue.

Motion capture through videography is a useful approach for evaluating the motor behaviors of humans and other species. Traditionally, motion capture has been performed by placing tracking markers on the target regions of the subject (Bernstein, 1967; Winter, 2009; Vargas-Irwin et al., 2010; Wenger et al., 2014; Maghsoudi et al., 2017). However, applying this approach to tongues present technical problems because tracking markers set on wet tongue regions can easily be displaced during tasks involving tongue movements. Moreover, using tracking markers pose risks in patients with tongue sensorimotor impairment as they may accidentally swallow the tracking markers. Regarding its clinical application, while setting objects on the tongue, it is important to reduce the risk of infections such as COVID-19 to the experimenter via the saliva.

Recently, Mathis et al. (2018) reported significant progress with the use of “DeepLabCut.” They implemented a systematic method to estimate the tracks of markerless movements. They successfully demonstrated that a small number of training images (~200) was sufficient to train this network with human-level labeling accuracy. This is possible because of transfer learning; the feature detectors are based on extremely deep neural networks, which were pretrained on ImageNet (He et al., 2016). Thus, this method involves the use of transfer learning techniques with deep neural networks, and yields outstanding results with minimal training data. This deep learning-assisted motion tracking system with DeepLabCut is useful for the application of tongue CKC because it does not use any recording device or tracking marker on the tongue, thereby eliminating the previously mentioned disadvantages of magnetic device noise, marker displacement, and additional risks of accidental aspiration and infection.

Herein, we introduce a novel approach that utilizes the CKC between the MEG and movement signals of the tongue during rhythmic tongue movements based on a deep learning-assisted motion capture system. Our main hypothesis is that the source locations for the tongue CKC differs from those of the finger CKC using the deep learning-assisted motion tracking system. In addition, to confirm the hypothesis that the CKC using the deep learning-assisted motion tracking system is reliable, we validate this CKC approach by comparing the CKC of fingers using motion capture with the CKC using ACC.

## 2. Materials and Methods

### 2.1 Subjects

Twelve healthy volunteers (10 men, 2 women; aged 21–35 y; mean age = 25.0 y) were examined. The participants were right-handed, as determined by the Edinburgh Handedness Inventory (Oldfield, 1971). None of the subjects had a history of neurological or psychiatric disorders. All the participants provided written informed consent in accordance with the ethical standards stated in the Declaration of Helsinki, and the study was approved by the local ethics and safety committees at Osaka University Hospital (No. 16469-2) and the Center for Information and Neural Networks (CiNet) at the National Institute of Information and Communications Technology (No. 1910280040).

### 2.2 Movement Tasks of Tongue and Fingers

The subjects were asked to perform constant, rhythmic side-to-side tongue movements with a slightly opened mouth for at least 3 min in two or three sessions (60–90 s each), separated by 30-s rest periods. They were asked to avoid drastic tongue movements to reduce the effects of touch sensations from the orofacial regions during tongue movement. They were also requested to relax the other orofacial parts during these tasks.

In the control task, the subjects were asked to make constant, rhythmic up-and-down movements of the right index finger over a table for at least 3 min in two sessions (90 s each), separated by a resting period of 30 s. During the resting periods, subjects were permitted to relax their orofacial muscles and swallow the saliva.

We attempted to observe the rhythmic movements of the right index finger in all twelve subjects (right finger condition). Four subjects (Subject 2, 3, 6, 12) performed rhythmical movements for both conditions (right and bilateral finger conditions) in a randomized order. The subjects were asked not to touch the table or other fingers during the finger movement tasks.

During the tongue and finger movement tasks, the participants were directed to fixate their gaze at a point on the wall in a magnetically shielded room to avoid any effects of eye movement or visual perception.

### 2.3 Recordings

#### 2.3.1 MEG and ACC recording

Cortical activity was recorded by CiNet using a whole-head MEG system with 360 channels (204 planar gradiometers, 102 magnetometers, and 54 axial gradiometers) (Neuromag® 360, Elekta, Helsinki, Finland). Planar gradiometers with 204 channels were used for the analysis. The position of the subject’s head inside the MEG helmet was continuously monitored by supplying a current to four coils fixed to the scalp for tracking head movements. An electromagnetic tracker was used to fix the coils according to the anatomical fiducials (Fastrak, Polhemus, Colchester, VT). The participants were seated in an upright position in the magnetically shielded room. To monitor the movements of the right index finger, a three-axis ACC (KXM52-1050, Kionix, Ithaca, NY, USA) was attached to the nail of the right index finger. The ACC cables were fixed to the hand and table using tape to prevent the generation of noise. The MEG and ACC signals were recorded with a passband at 0.03–330 Hz, and the signals were sampled at 1 kHz.

#### 2.3.2 Video and MRI recording

The movements of each target region (the tongue and index fingers) were video-recorded simultaneously throughout the MEG recording at 120 fps with a resolution of 1280 × 720 pixels, using a camera (DMC-FZ200, Panasonic, Osaka, Japan). To obtain a frontal view of each target region, the camera was positioned in front of the MEG gantry at a distance of 1.5 m. To record the finger and tongue movements, the zoom function of the camera was used to record the images of both hands—including the index fingers—and the lower part of the orofacial region (from neck to nasion). To match the onset time between the MEG and movement signals with motion capture analysis, the MEG system included a light-emitting diode (LED) that was strobed five times at 1 Hz before and after each movement task and was captured in the video images. To determine the brain anatomy of each subject, three-dimensional T1 magnetic resonance images (MRIs) were acquired using a 3T MRI scanner (Siemens MAGNETOM Trio or Vida, Siemens, Munich, Germany).

### 2.4 Data Analysis

#### 2.4.1 Movement signals with the motion capture system

The movements of the tongue and fingers were analyzed offline via deep learning-assisted motion capture with videography using the open-source toolbox, DeepLabCut (Mathis et al., 2018) (https://github.com/AlexEMG/DeepLabCut). The image resolution was changed to 960 × 540 pixels. For motion tracking, we extracted 100–150 random, distinct frames from the videos for each movement task. We cropped the frames such that the target regions were clearly visible and manually labeled the tip of the tongue/finger in each extracted frame. The system was then trained using a deep neural network architecture to predict the target regions based on the corresponding images. Different networks were trained for each target region in 100,000–200,000 iterations as the loss relatively flattened (Mathis et al., 2018; Nath et al., 2019). The trained networks could track the locations of the target regions in the full sets of video segments (Supplementary Videos 1 and 2). The labeled *x-*axis (i.e. left-right) and *y-*axis (i.e. bottom-top) positions of the pixels in each frame were stored and exported in CSV format for subsequent analysis using MATLAB (The MathWorks, Natick, Massachusetts, USA). The Euclidian norm of the two orthogonal (*x*- and *y*-axes) signals with baseline correction was used as the movement signal for motion capture.

#### 2.4.2 Coherence between MEG and movement signals

The raw MEG signals were spatially filtered offline with the temporal extension of the signal space separation method (Taulu and Simola, 2006; Taulu and Hari, 2009) using MaxFilter (version 2.2.12, Elekta Neuromag, Finland). The MEG and ACC signals were adjusted by down-sampling to 500 Hz. The movement signals were adjusted by up-sampling with the motion capture system to match the MEG signals at 500 Hz. LED flashes were applied to the images for correction between the MEG and movement signals with motion capture.

The coherence spectra between the MEG and rectified movement signals with motion capture were calculated using the method proposed by Welch (1967) for the estimation of spectral density, where half-overlapping samples, a frequency resolution of 0.5 Hz, and a Hanning window were used. The following equation was used to determine the coherence (Coh*xy*).

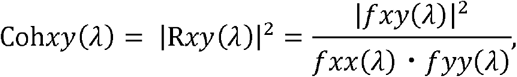

where *fxx*(*λ*) and *fyy*(*λ*) respectively denote the values of the auto-spectra of the MEG signals and rectified movement signals with motion capture for a given frequency, λ, and *fxy*(*λ*) represents the cross-spectrum between *fxx*(*λ*) and *fyy*(*λ*). We used the position data as movement signals for the CKC analysis with capture motion since the mean CKC value is within 5% error among approaches using position, velocity, and acceleration (Supplementary Table). The coherence spectra between the MEG and Euclidian norm of the three orthogonal ACC signals (*x*-axis (i.e. left-right), *y*-axis (i.e. bottom-top), *z*-axis (i.e. near-far)) from right index finger were also calculated.

We checked the epochs comprising artifacts related to unintended orofacial muscle movements such as coughing, which were distinguished through visual inspection. 96.83 ± 1.79 (mean ± standard error of the mean (SEM)) (ranging from 88 to 107 (*n* = 12)) samples were obtained for the tongue CKC. The epochs for the finger CKC included 96.42 ± 1.52 (ranging from 87 to 106 (*n* = 12)) samples for the right finger condition and 105.00 ± 3.24 (ranging from 98 to 111 (*n* = 4)) samples for the bilateral finger condition. According to the method proposed by Rosenberg et al. (1989), all coherence values above *Z* were considered to be significant at *p* < 0.01, where *Z* = 1−0.01^(1/*L*−1)^ and *L* denotes the total number of samples for the auto- and cross-spectrum analyses.

The cross-correlogram in the time domain was calculated by applying an inverse Fourier transformation to the averaged cross-spectra for the tongue CKC and right finger CKC with motion capture. The cross-correlogram underwent bandpass filtering at 1–45 Hz. Isocontour maps were constructed at the time points at which the peaks of the cross-correlogram were observed. The sources of the oscillatory MEG signals were modeled as equivalent current dipoles (ECDs). To estimate the ECD locations, the spherical head model was adopted; the center of this model was consistent with the local curvature of the brain surface of an individual, as determined by the MRI (Sarvas, 1987). Only the ECDs with a goodness-of-fit value of at least 85% were accepted. One subject (Subject 11) was excluded from the ECD analysis of the tongue CKC due to an insufficient goodness-of-fit criterion.

### 2.5 Statistical analysis

The data are expressed as the mean ± SEM. An arc hyperbolic tangent transformation was used to normalize the values of the coherence to ensure that the variance was stabilized (Halliday et al., 1995). The values of the CKC of the tongue were analyzed between the left and right hemispheres using paired *t*-tests. The statistical significance level was set to *p* < 0.05. The ECD locations over the left hemisphere along each axis (*x*-, *y*-, and *z*-axes) were analyzed between the tongue CKC and right finger CKC using paired *t*-tests with Bonferroni correction. The corrected *p* value with Bonferroni correction was set to *p* < 0.0167 (0.05/3). The *x*-axis intersected the preauricular points from left to right; the *y*-axis intersected the nasion; the *z*-axis was perpendicular to the plane determined by the *x*- and *y*-axes.

## 3. Results

Figures 1A and B depict representative raw data and power spectra of the movement signals with motion capture and the ACC, respectively, for the right finger condition of Subject 2. Cyclic rhythms were observed at a specific frequency band of the finger movements for both motion capture and the ACC (Fig. 1A). The peak of the power spectra of movement signals with both motion capture and the ACC exhibited the same frequency band of movement rhythms, at 3.3 Hz (indicated by arrows) (Fig. 1B). The peak CKC of the right finger was observed over the contralateral hemisphere at 7.0 Hz with both motion capture (CKC value = 0.61) and the ACC (CKC value = 0.60), around the harmonic frequency band of finger movements (Fig. 1C). The peak CKC of the tongue was observed over the left hemisphere (CKC value: 0.43) and right hemisphere (CKC value: 0.46) at 3.3 Hz, around the harmonic frequency band of tongue movements (Fig. 2A[1,2]).

**Fig. 1.**
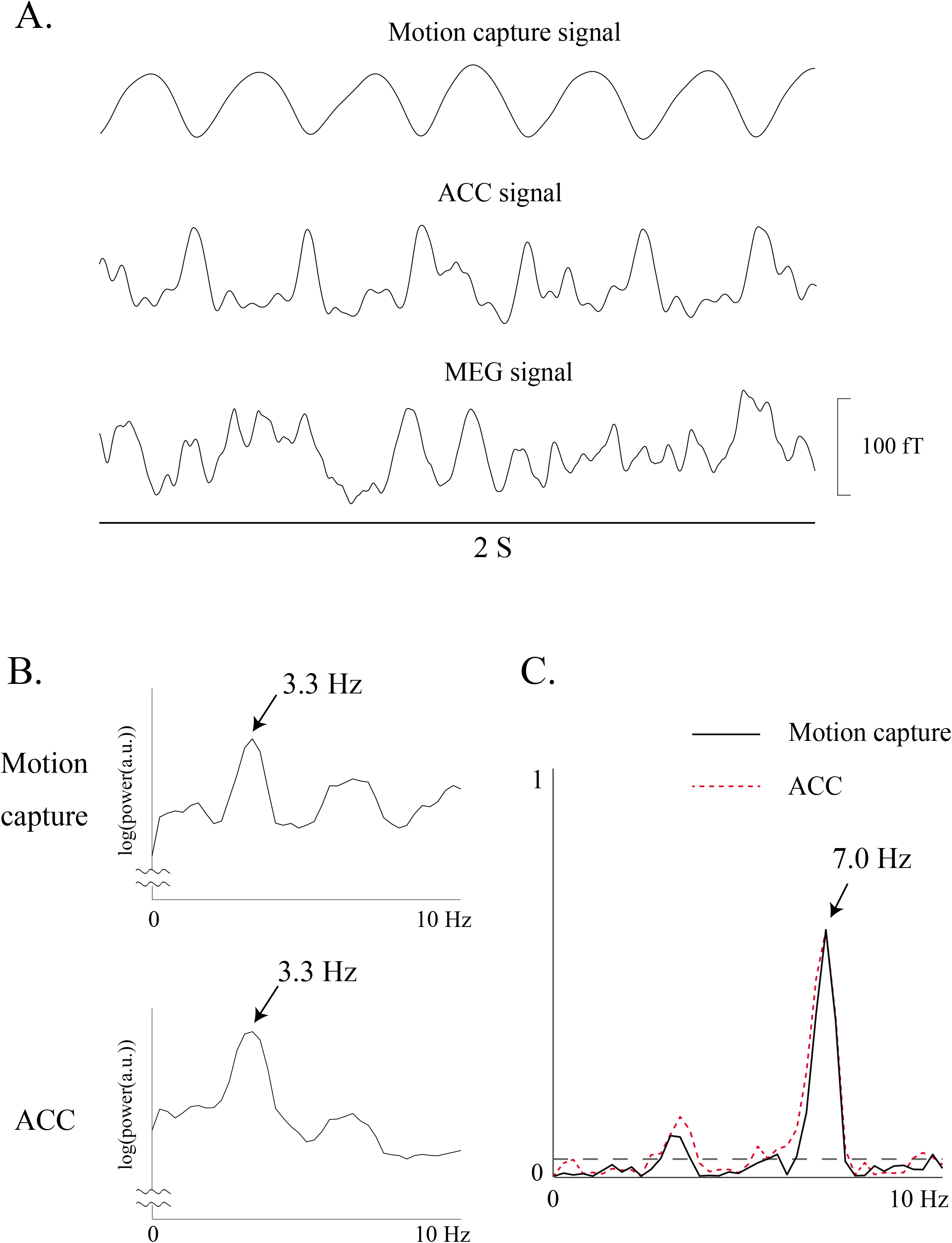
A. Raw data of movement signals obtained through motion capture and an accelerometer (ACC), and magnetoencephalographic (MEG) signal from the contralateral (left) Rolandic sensor for the right finger movement condition of a single participant (Subject 2). Cyclical rhythms are observed at a specific frequency band of finger movements using both the motion capture and ACC. B. Power spectra of movement signals obtained through motion capture and the ACC for the right finger movement condition of a single participant (Subject 2). The scale of the *x*-axis is 10 Hz. Note that the peak frequency occurs in the same frequency band of finger movement, i.e., at 3.3 Hz, in both the motion capture and ACC results (indicated by arrows). C. Corticokinematic coherence (CKC) waveform from a representative channel over the contralateral hemisphere for the right finger movement condition of a single participant (Subject 2) using motion capture and the ACC. The scale of the *x*-axis is 10 Hz. The horizontal dashed line indicates a significance level of 99%. The CKC peak is observed at 7.0 Hz in the motion capture (CKC value: 0.61) and ACC (CKC value: 0.60) results around the harmonic frequency band of the finger movements.

**Fig. 2.**
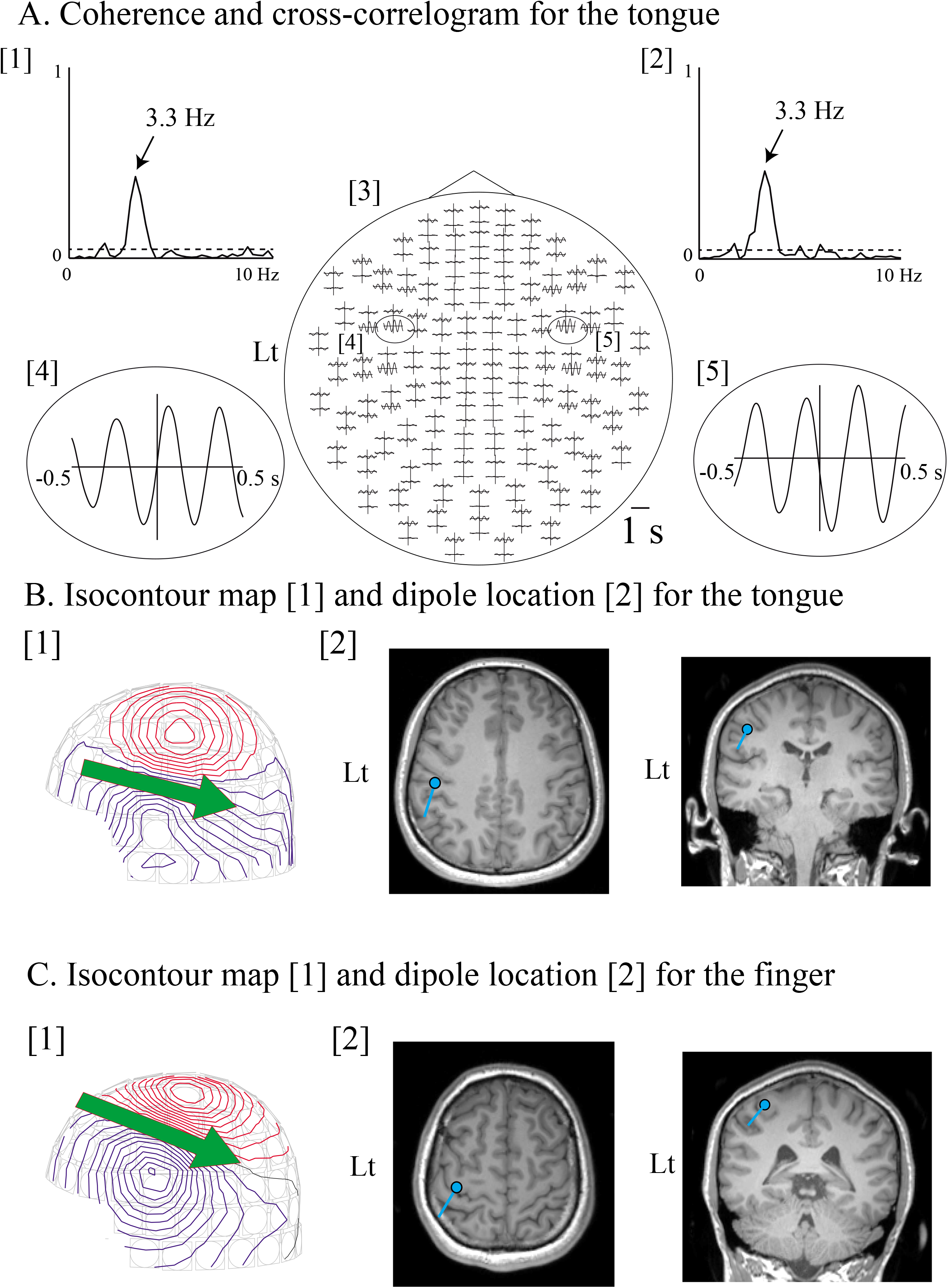
A. [1, 2] Corticokinematic coherence (CKC) waveform for the tongue from a representative channel over the left [1] and right [2] hemispheres of a single participant (subject 1). The scale of the *x*-axis is 10 Hz. The horizontal dashed line indicates a significance level of 99%. The CKC peak is observed at 3.3 Hz in the left hemisphere (CKC value: 0.43) and right hemisphere (CKC value: 0.46). [3-5] Spatial distribution of the 1-s-long cross-correlogram for the tongue of a single participant (subject 1). The largest peaks of the cross-correlogram occurred in the Rolandic sensors of the left [4] and right [5] hemispheres for the tongue CKC. B. Isocontour maps and dipole locations for the tongue (B) and finger (C) of Subject 1. The time points that showed the cross-correlation peaks were used to obtain the contour map. The incoming and outgoing magnetic fluxes are denoted by the blue and red lines, respectively (B[1], C[1]). The green arrows denote the directions and locations of the equivalent current dipoles (ECDs), which were projected onto the surface of the skull. The arrowheads indicate the negative poles of the ECDs. The ECDs (blue dots) of the tongue (B[2]) and finger (C[2]) are superimposed on magnetic resonance image slices of the participant. The directions of the blue lines represent the negative poles of the ECDs. Both ECDs are located at the central sulcus (B[2], C[2]). The locations of the ECDs of the tongue are estimated to be more lateral, anterior, and inferior to those of the finger. Lt: Left side.

For the right finger condition, the peak frequencies of the power spectrum of the movement signals were the same, at 1.8–3.8 Hz for both motion capture and the ACC (Table 1). The coherence spectra exhibited significant peaks (*p* < 0.01) over the contralateral hemisphere at 2.0–7.0 Hz and 2.0–7.0 Hz with motion capture and the ACC, respectively, corresponding to the frequencies of finger movements or their harmonics in all 12 subjects (Table 1). The CKC value with motion capture (mean, 0.433) was compared with that of CKC with the ACC (mean, 0.510), achieving a similarity of 84.9% (Table 1). For the bilateral finger condition, the CKC also exhibited peaks for each side of the finger in all 4 subjects (Table 2).

**Table 1.** Peak frequency and values of CKC of the fingers—right finger conditions

| Sub | Peak frequency (Hz) |  |  |  | CKC value |  |
| --- | --- | --- | --- | --- | --- | --- |
|  | Movement signal |  | CKC |  | ACC | Motion capture |
|  | ACC | Motion capture | ACC | Motion capture |  |  |
| 1 | 1.8 | 1.8 | 3.3 | 3.3 | 0.80 | 0.69 |
| 2 | 3.3 | 3.3 | 7.0 | 7.0 | 0.60 | 0.61 |
| 3 | 2.0 | 2.0 | 4.0 | 4.0 | 0.47 | 0.44 |
| 4 | 3.8 | 3.8 | 3.3 | 3.3 | 0.41 | 0.32 |
| 5 | 2.0 | 2.0 | 4.0 | 3.5 | 0.44 | 0.55 |
| 6 | 1.8 | 1.8 | 3.3 | 3.3 | 0.65 | 0.49 |
| 7 | 1.8 | 1.8 | 3.8 | 3.8 | 0.35 | 0.32 |
| 8 | 2.8 | 2.8 | 3.0 | 5.5 | 0.33 | 0.34 |
| 9 | 2.0 | 2.0 | 4.0 | 3.8 | 0.56 | 0.47 |
| 10 | 2.0 | 2.0 | 2.0 | 2.0 | 0.66 | 0.55 |
| 11 | 2.5 | 2.5 | 5.0 | 5.3 | 0.55 | 0.26 |
| 12 | 2 | 2 | 2 | 2 | 0.29 | 0.19 |
| Ave | 2.32 | 2.32 | 3.56 | 3.49 | 0.510 | 0.433 |
| Min | 1.8 | 1.8 | 2.0 | 2.0 | 0.29 | 0.19 |
| Max | 3.8 | 3.8 | 7.0 | 7.0 | 0.80 | 0.69 |
| SEM | 0.19 | 0.19 | 0.41 | 0.43 | 0.044 | 0.043 |

ACC: accelerometer; Ave: average; CKC: corticokinematic coherence; Max: maximum; Min: minimum; Movement signal: power spectrum of the movement signal; SEM: standard error of the mean; Sub: subject number.

**Table 2.** Peak frequency and values of CKC of the fingers—bilateral finger conditions

| Sub | Peak frequency (Hz) |  |  |  |  |  | CKC value |  |  |
| --- | --- | --- | --- | --- | --- | --- | --- | --- | --- |
|  | Movement signal |  |  | CKC |  |  |  |  |  |
|  | ACC | Motion capture |  | ACC | Motion capture |  | ACC | Motion capture |  |
|  | Rt | Rt | Lt | Rt | Rt | Lt | Rt | Rt | Lt |
| 2 | 3.0 | 3.0 | 3.0 | 6.0 | 6.0 | 3.3 | 0.48 | 0.22 | 0.22 |
| 3 | 2.3 | 2.3 | 2.3 | 2.3 | 2.3 | 2.3 | 0.69 | 0.49 | 0.38 |
| 6 | 2.0 | 2.0 | 2.0 | 2.0 | 2.0 | 4.3 | 0.36 | 0.26 | 0.20 |
| 12 | 2.5 | 2.5 | 2.5 | 5.0 | 2.5 | 2.8 | 0.36 | 0.24 | 0.22 |

ACC: accelerometer; CKC: corticokinematic coherence; Lt: left finger; Movement signal: power spectrum of the movement signal; Rt: right finger; Sub: subject number.

For the tongue movements, the peak frequencies of the power spectrum of the movement signals were detected at 1.3–3.3 Hz (Table 3). The CKC spectra for the tongue showed significant peaks (*p* < 0.01) at 2.5–5.3 Hz over the left hemisphere and at 2.5–6.0 Hz over the right hemisphere in all subjects, corresponding to the frequency of tongue movements or their harmonics (Table 3). The CKC values were not significantly different between the left (mean, 0.203) and right (mean, 0.188) hemispheres (*p* = 0.499) (Table 3).

**Table 3.**
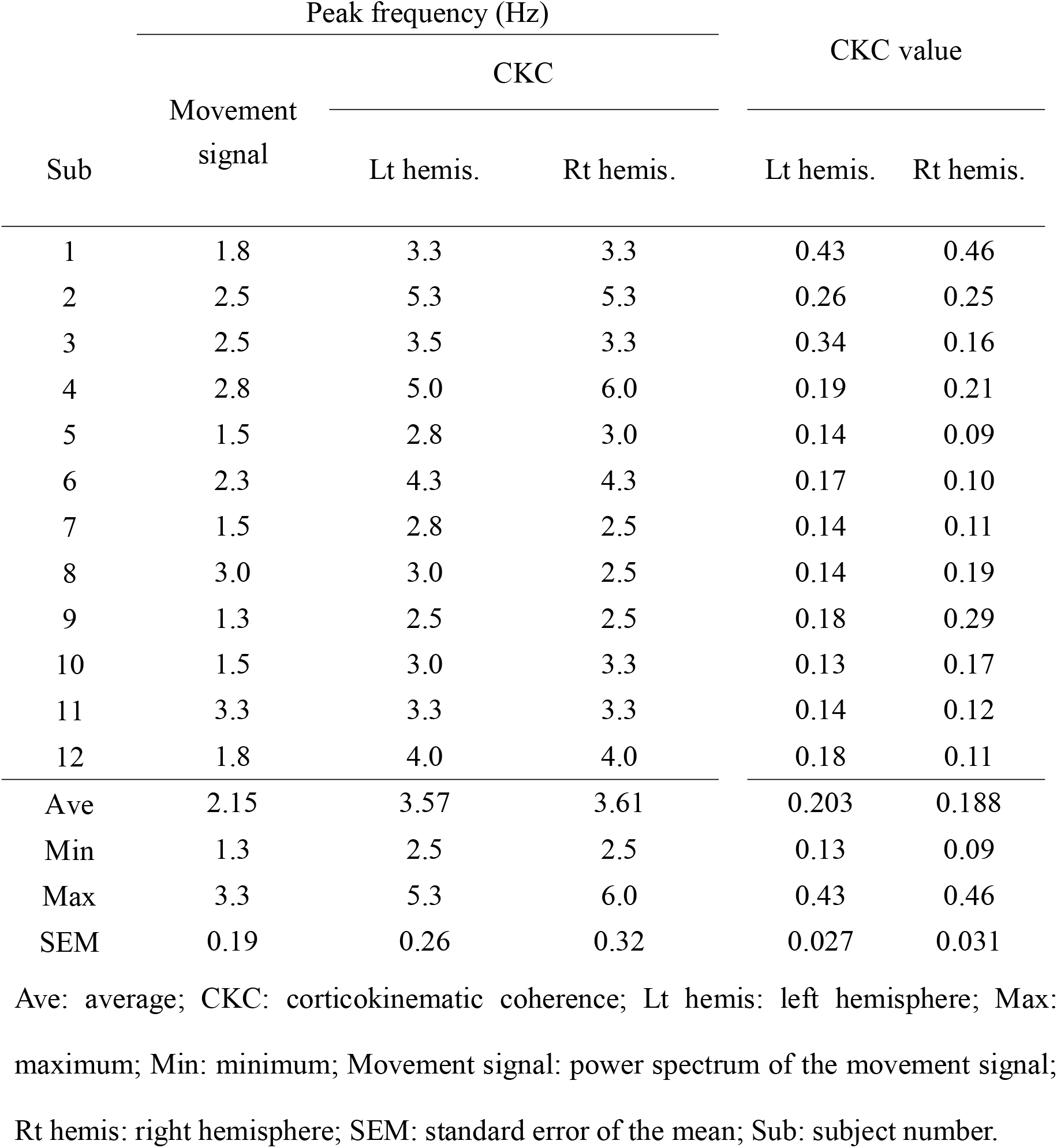
Peak frequency and values of CKC of the tongue

The spatial distributions of the cross-correlogram of the finger and tongue CKC showed peaks over the contralateral and bilateral hemispheres (Fig. 2A[3–5]), respectively. Dipolar field patterns, which were centered on the Rolandic sensors, were observed at the principal peaks of the cross-correlogram (Fig. 2B[1]). The sources for the tongue CKC were estimated to be over the left and right SM1 in 11 subjects, respectively (Fig. 2B[2]). The sources for the right finger CKC were located in the SM1 over the contralateral hemisphere in all of the 12 subjects (Fig. 2C[2]). The results of the paired *t*-test implied that the locations of the ECDs of the tongue were considerably lateral (mean = 13.99 mm; *p* < 0.001; paired *t*-test with Bonferroni correction) and inferior (mean = 20.78 mm; *p* < 0.001), but not anterior (mean = 5.15 mm; *p* = 0.029) to those of the finger (Fig. 3).

**Fig. 3.**
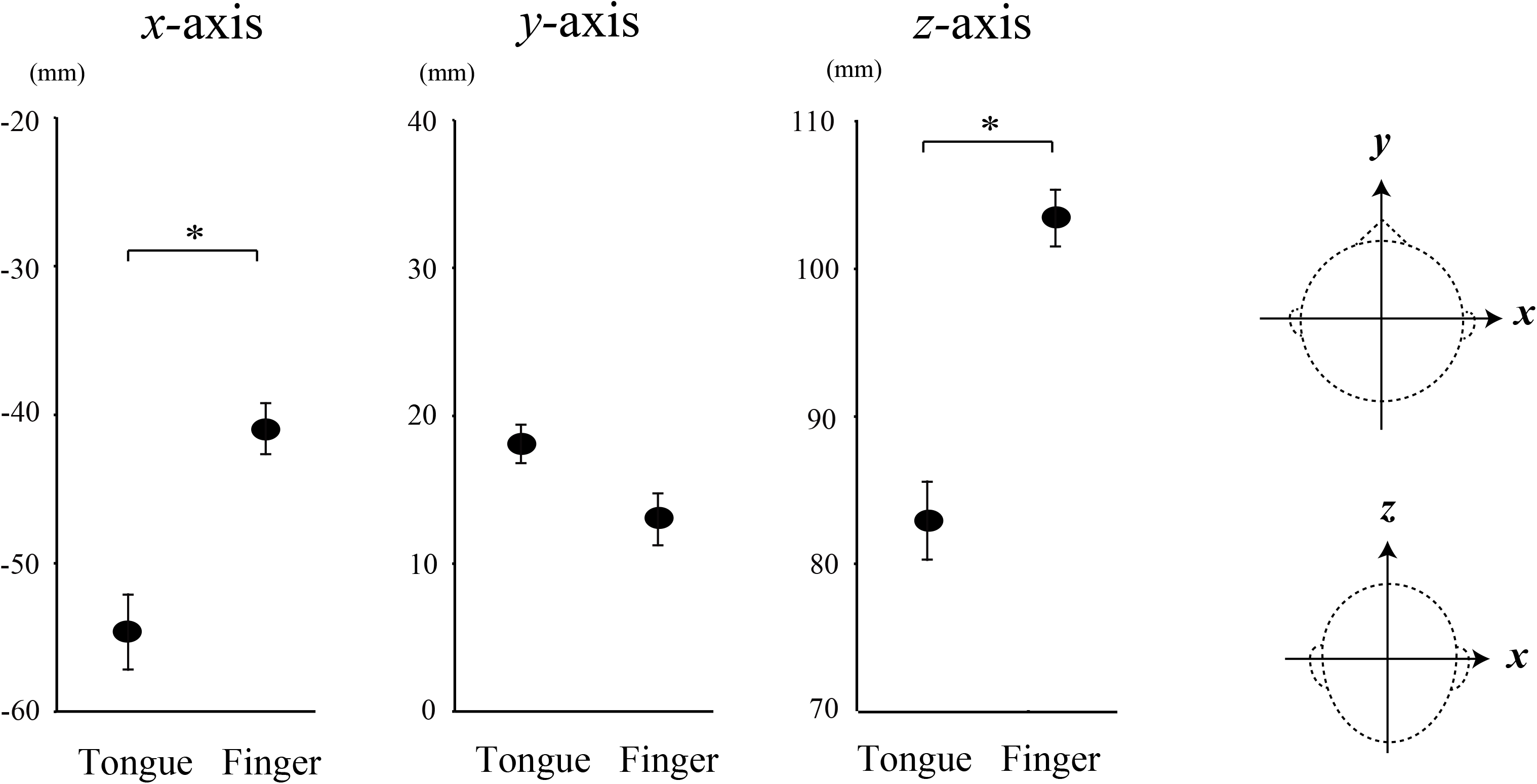
Average locations of the ECDs of the tongue and finger CKCs on the *x*-, *y*-, and *z*-axes, considering all participants. The data points represent the means ± SEM values. The locations of the ECDs of the tongue are considerably lateral and inferior to those of the finger. The *x*-axis intersects the preauricular points from left to right; the *y*-axis passes through the nasion; the *z*-axis is perpendicular to the plane determined by the *x*- and *y*-axes. Asterisks indicate statistically significant differences (*p* < 0.0167).

## 4. Discussion

Significant coherence between MEG and tongue movement signals was detected over the bilateral hemispheres using deep learning-assisted motion capture with videography. The sources of the coherence activity were detected in the bilateral SM1 of the tongue region, which were found to be considerably lateral and inferior to the finger SM1, corresponding to the classical homunculus. These results suggest that the use of deep learning-assisted motion capture in CKC is a robust and useful approach for evaluating the functional localization of the tongue SM1.

The reliability of measuring CKC using motion capture is comparable to that of the conventional ACC-based CKC method (Bourguignon et al., 2011; 2012), as evidenced by the fact that the finger CKC value obtained using motion capture achieved a similarity of 84.9% when compared with the CKC value obtained using the ACC and the finger CKC value obtained using ACC. In addition, as the power spectrum of movement signals and CKC showed the same peak frequency bands between the motion capture and ACC for all subjects during the finger movement tasks, determining the CKC with deep learning-assisted motion capture was found to be reliable.

Previous studies involving non-human primates have revealed that several movement parameters, such as position, rotation, direction, and movement velocity, are encoded in the SM1, as determined using the recordings of a single neuron, local field potential, and multi-unit activity (Ashe and Georgopoulos, 1994; Caminiti et al., 1990; Carmena et al., 2003; Mehring et al., 2003; Moran and Schwartz, 1999; Reina et al., 2001). MEG studies involving humans have also revealed the significance of the SM1 cortex oscillations for encoding the parameters of voluntary movements, such as velocity (Jerbi et al., 2007) and acceleration (Bourguignon et al., 2011; 2012). When studying CKC with motion capture, we evaluated the movement parameters of the target positions of pixels in each image with videography by using a deep learning-assisted motion capture system, since the CKC value with motion capture is not significantly different among approaches using position, velocity, and acceleration (Supplementary Table).

Recently, Bourguignon et al. (2019) reported that using two different approaches showed interactions between central and peripheral body parts during motor executions; i.e. CKC and cortico-muscular coherence (CMC) occurs by different mechanisms. CKC, which is coherent with the movement frequency and its harmonics, is mainly related to proprioceptive afferent signals. CMC, which mainly occurs at beta frequency bands during weak muscle contraction, is mainly driven by mu-rhythm-specific neural modulations in efferent signals. Bourguignon et al. (2019) also reported that the values of CKC during rhythmic finger movements were substantially higher and easier to detect than those of CMC during isometric finger movements (Brown et al., 1998; Conway et al., 1995; Farmer et al., 1993; Gross et al., 2000; Halliday et al., 1998; Kilner et al., 1999, 2004; Mima et al., 1999; Salenius et al., 1997). Because a recording time of at least 10 min was required for the CMC of the tongue in previous studies (Maezawa et al., 2014; 2016), the proposed motion capture approach offers the advantage of a short recording time—approximately 3 min for the CKC of the tongue. The CKC of the tongue with motion capture also has a technical advantage of enabling free movement because no objects, such as an ACC, electromyography (EMG) electrodes, or tracking markers, are placed on the tongue. When objects are placed on the tongue, they disturb the execution of smooth movement tasks. For example, for the tongue CMC recording, it is sometimes technically challenging to set the EMG electrodes on narrow and wet tongue regions because placing electrodes on the tongue can induce uncomfortable feelings in subjects, resulting in a vomiting reflex. Moreover, because no objects are used on the tongue in this CKC method, the risk of an object being swallowed during a tongue movement task is eliminated. In clinical applications for patients with sensorimotor disorders of the tongue, patients sometimes face difficulties performing smooth tongue movements and are easily fatigued by movement tasks. Therefore, the short recording time of the tongue CKC technique provides an advantage over the conventional CKC and CMC methods that use ACC devices or EMG electrodes. In a recent clinical setting, Marty et al. (2019) reported that utilization of the finger CKC is a useful approach for patients with impairment of spinocortical proprioceptive pathways in Friedreich ataxia. As oropharyngeal dysphagia and/or speech disorders are also commonly present in individuals with Friedreich ataxia and worsens with disease duration and severity (Keage et al., 2017), the CKC approach of the tongue might provide electrophysiological evidence for proprioceptive impairment of corticobulbar proprioceptive pathways.

Damage to the cortical areas representing sensorimotor function of the extremities and language function causes severe dysfunction and seriously decreases the quality of life. Thus, cortical localization of these functions has received much attention for the presurgical evaluation of neurosurgical procedures. In contrast, cortical localization of functions relating to the tongue and other orofacial regions has been relatively undervalued. This is because the cortical representation of orofacial motor function is bilateral, and thus damage to the orofacial SM1 does not apparently induce severe dysfunctions unless the damage is bilateral as well (Cukiert et al., 2001; Lehman et al., 1994). However, dysfunctions in critical orofacial motor functions may still result from damage to the orofacial SM1, severely reducing the quality of life. For example, dysfunctions in critical tongue motor functions can cause dysphagia and silent aspiration (Meadows, 1973; Horner and Massey, 1988; Robbins et al., 1993). In addition, damage to the orofacial SM1 may cause a cosmetically conspicuous imbalance of facial expression between the left and right sides of the face (Lehman et al., 1994). Because this unbalanced facial expression is easily recognized in daily communication, the problem should be considered as a target for improvement. Thus, more attention should be paid to preserving motor functions of the tongue and other orofacial regions during neurosurgical operations. Here, the CKC technique may be helpful in evaluating SM1 localization of the orofacial regions in patients with brain lesions observed around the central sulcus.

Previous studies have shown that the finger CKC mainly reflects the proprioceptive input into the contralateral SM1 (Piitulainen et al., 2013; Bourguignon et al., 2015), which corresponds to the timing of the strongest deflection of the cortical MEFs associated with self-paced finger movements (Cheyne et al., 1997). Thus, it is likely that the cortical mechanisms of the CKC and MEFs are closely related; therefore, it is reasonable that the tongue CKC was detected over both SM1s without hemispheric dominance—similar to the MEF results obtained in the bilateral SM1 associated with self-paced tongue protrusion tasks with intervals of approximately 10 s (Maezawa et al., 2017).

Previous studies have reported that the CMC for the tongue was detected at 2–10 Hz, which may have been driven by proprioceptive afferents from the tongue muscles to the cortex—as well as the beta frequency band—during sustained tongue protrusion tasks (Maezawa et al., 2014; 2016). Because human tongue muscles are rich in muscle spindles (Cooper, 1953), it is reasonable that the tongue CKC may be related to the proprioceptive afferents from the tongue muscles associated with rhythmic tongue movements.

Ruspantini et al. (2012) reported that low oscillatory frequency, which is related to the proprioceptive afferent feedback obtained from the mouth muscles, might be necessary to generate the fine oral movements required to produce speech. Therefore, sensory feedback obtained by muscle spindles of the orofacial regions may contribute to excellent oral motor functions, including swallowing, speech, and mastication. CKC with motion capture has the advantage of being able to track the motions of multiple body parts, as the finger CKC for bilateral finger movements can be evaluated simultaneously. Thus, in the future, CKC with motion capture might be useful for elucidating the cortical mechanisms that enable swallowing and speech through evaluation of the synchronization of signals between the MEG and movements of multiple orofacial regions.

The occurrence of synchronous head movements corresponding to rhythmic tongue movements may yield coherent artifacts in the cross-correlogram. This feature represents a potential limitation of the tongue CKC during repetitive tongue movements, similar to the limitations related to the finger CKC mentioned in previous studies (Bourguignon et al., 2011; 2019). In clinical applications of the tongue CKC, the appearance of artifacts related to head movements must be addressed in patients who struggle to perform repetitive movements. Another potential limitation is the effect of touch sensations from the tongue and other orofacial regions, such as the buccal and lip, during tongue movement tasks. Because CKC appears to be primarily driven by proprioceptive feedback with no significant evidence of any effect due to cutaneous input (Piitulainen et al., 2013; 2015), touch sensations might not have been a severe problem in the present study. Further studies are required to analyze the effects of touch sensations from orofacial regions on the tongue CKC during tongue movement tasks. We applied single dipole fitting analysis for the source localization for clinical application, as dipole fitting is useful for evaluating the somatotopic localization in a pre-neurosurgical situation. However, it is also useful to reveal the distribution of cortical activity based on the distributed source modelling from the systematic and physiological point of view. Further studies are needed to reveal the cortical mechanisms of tongue movements using distributed source modelling analysis.

In conclusion, the use of CKC together with deep learning-assisted motion capture is a robust and useful approach for evaluating the functional localization of the SM1 of the tongue; it is a magnetic, noise-free, movement-free, and risk-free approach because no recording devices are placed on the tongue.

## Abbreviations

ACC: Accelerometer
CKC: Corticokinematic coherence
ECD: Equivalent current dipole
EMG: Electromyography
LED: light-emitting diode
MEG: Magnetoencephalography
MEF: Movement evoked field
MRI: Magnetic resonance image
SM1: primary sensorimotor cortex
SEM: Standard error of the mean.

## Acknowledgments

We thank Dr. Takafumi Suzuki, Dr. Takeshi Nogai and Dr. Asuka Otsuka (Center for Information and Neural Networks (CiNet), National Institute of Information and Communications Technology, and Osaka University) for supporting our MEG recording.

## Supplementary Video 1

Sample video of pose estimation of the tongue during the tongue movement task. The solid blue circles were identified using the learning program with DeepLabCut. The movie is slowed to a quarter of the real-time speed.

## Supplementary Video 2

Movement task of the fingers in the bilateral finger condition. The solid blue (right finger) and red (left finger) circles were identified using the learning program with DeepLabCut. The movie is slowed to a quarter of the real-time speed.

