## Supplementary_Table for "Functional cortical localization of the tongue using corticokinematic coherence with a deep learning-assisted motion capture system"

|  | Peak frequency (Hz) | | | | | |  | CKC value | | |
| --- | --- | --- | --- | --- | --- | --- | --- | --- | --- | --- |
|  | Movement signal | | | CKC | | |  |  |  |  |
| Sub | P | V | A | P | V | A |  | P | V | A |
| 1 | 1.8 | 3.3 | 3.3 | 3 | 3 | 3 |  | 0.69 | 0.69 | 0.71 |
| 2 | 3.3 | 3.3 | 3.3 | 7 | 7 | 7 |  | 0.61 | 0.59 | 0.45 |
| 3 | 2.0 | 2.0 | 2.0 | 4.0 | 4.0 | 4.0 |  | 0.44 | 0.44 | 0.45 |
| 4 | 3.8 | 3.8 | 3.8 | 3.3 | 3.3 | 3.3 |  | 0.32 | 0.31 | 0.31 |
| 5 | 2.0 | 2.0 | 2.0 | 3.5 | 3.5 | 3.5 |  | 0.55 | 0.53 | 0.44 |
| 6 | 1.8 | 1.8 | 1.8 | 3.3 | 3.3 | 3.33 |  | 0.49 | 0.41 | 0.45 |
| 7 | 1.8 | 1.8 | 1.8 | 3.8 | 3.8 | 3.8 |  | 0.32 | 0.31 | 0.30 |
| 8 | 2.8 | 2.8 | 2.8 | 5.5 | 5.5 | 5.5 |  | 0.34 | 0.34 | 0.34 |
| 9 | 2.0 | 2.0 | 2.0 | 2.0 | 2.0 | 2.0 |  | 0.47 | 0.43 | 0.44 |
| 10 | 2.0 | 2.0 | 2.0 | 2.0 | 2.0 | 2.0 |  | 0.55 | 0.56 | 0.56 |
| 11 | 2.5 | 2.5 | 2.5 | 2.5 | 2.5 | 2.5 |  | 0.26 | 0.28 | 0.29 |
| 12 | 2.0 | 2.0 | 2.0 | 2.0 | 2.0 | 2.0 |  | 0.19 | 0.20 | 0.23 |
| Ave | 2.44 | 2.44 | 2.44 | 3.49 | 3.49 | 3.49 |  | 0.433 | 0.424 | 0.413 |
| SEM | 0.20 | 0.20 | 0.20 | 0.43 | 0.43 | 0.43 |  | 0.043 | 0.042 | 0.038 |

P: Position; V: Velocity; A : Acceleration

To assess the validation of the corticokinematic coherence (CKC) with motion capture system using position data, we compared the peak frequencies of power spectrum for motion signals and CKC among approaches using position, velocity and acceleration in the right finger conditions. We also compared the CKC value among approaches using position, velocity and acceleration. The power of motion signals and CKC shows the same peak frequency bands among position, velocity and acceleration, respectively, in all of the subjects. Moreover, as the CKC value obtained from the position (mean: 0.433) reached a similarity of 100.21% and 100.46% when compared with the CKC value from the velocity (mean: 0.424) and acceleration (mean: 0.414), the CKC with capture motion system using position data was found to be a reliable and robust method.
